# The nasopalatine ducts of the mouse conserve a functional role in pheromone signaling

**DOI:** 10.1101/757930

**Authors:** Dana Rubi Levy, Yizhak Sofer, Vlad Brumfeld, Noga Zilkha, Tali Kimchi

## Abstract

Social communication in most mammals is mediated by chemosignals, collected by active sniffing and detected mainly by the vomeronasal organ (VNO). In reptiles, however, chemosignals are delivered to the VNO through the oral cavity via the nasopalatine ducts (NPDs) – two direct passageways connecting the nasal and the oral cavities. While the structure of the NPDs is highly conserved across terrestrial vertebrate, it is unclear whether they retain any functional role in mammalian chemosignaling. Here we assess the role of the mouse NPDs in VNO function and associated behavioral responses. By reconstructing the 3D morphological architecture of the mouse snout using micro CT, we identify a net of micro-tunnels forming a direct passageway connecting the NPDs to the nasal cavity and the vomeronasal organ. We further demonstrate that physical obstruction of the NPDs destructs VNO clearance, and reduces chemosignaling-evoked neuronal activation in the medial amygdala. Obstruction of the NPDs also impaired the innate male preference for female chemosignals as well as social approach behavior, indicating the crucial role of the murine nasopalatine ducts in pheromone sensing.

## Introduction

It is well accepted that the vomeronasal organ (VNO) plays a key role in perceiving sex-specific and species-specific chemical signals (Bear et al 2016, Dulac & Torello 2003, Halpern & Martinez-Marcos 2003, Isogai et al 2011, Keverne 1999, Kimchi et al 2007, Marom et al 2019, Stowers et al 2002). In most animal species, VNO-mediated signals regulate a variety of innate behaviors crucial for the survival of the individual and the species (Baum & Kelliher 2009, Bielsky & Young 2004, Brennan & Zufall 2006, Chamero et al 2007, Grinevich & Stoop 2018, Haga et al 2010, Papes et al 2010). An accumulating body of evidence reported that the mammalian VNO opens into the nasal cavity through a sole opening in its anterior end. Thus, chemosignals are thought to reach the VNO through the nostrils, by active sniffing (Dulac & Torello 2003, Vaccarezza et al 1981), while the VNO itself serves as an active pump to guide soluble molecules into its epithelium (Eccles 1982, Meredith 1994). Yet, a full characterization of how chemosignals reach the VNO is still missing.

In the chemosignaling system of reptiles, molecules are known to reach the VNO not through the nose, but rather through the mouth - via two fine tubular structures termed the nasopalatine ducts (NPDs). These ducts create a direct and continuous passage between the nasal and oral cavities, and present high evolutionary conservation across terrestrial vertebrate species (Shimp et al 2003, Wohrmann-Repenning 1980, Wohrmann-Repenning 1993). Felids and ungulates, for example, utilize the NPDs for pheromone transfer to the VNO by performing the distinct “flehmen” behavior, in which an animal curls back its upper lip exposing its front teeth and inhales, with the nostrils usually closed (Stahlbaum & Houpt 1989). Despite the fact that the NPDs are clearly present in most mammalian species (Jacob et al 2000, Shimp et al 2003), their role in mammalian chemosignaling and related behaviors has been usually overlooked, and the few studies which explored their functions yielded inconclusive results (Mackaysim & Rose 1986, Meredith 1991). Furthermore, flehmen behavior was not observed in small rodents such as mice and rats. Consequently, chemosignaling in mammals is still generally considered a “nasal” property, and there is uncertainty as to whether, and how, the NPDs are involved in this process.

In the present study, we explored the mechanism that enables influx of chemosignals to the mammalian VNO while looking beyond the common roles of the olfactory systems, and focusing on the importance of the nasopalatine ducts and the oral cavity. Considering their location and evolutionary functions, we hypothesized that the nasopalatine ducts are in fact an essential component of the mammalian chemosignaling system and facilitate substance flow to this organ.

Using high-resolution micro-computerized tomography (CT), together with *in-vivo* florescent tracing, we explored the flow path of liquid-borne compounds to the vomeronasal neuroepithelium. By permanently obstructing the oral openings of the NPDs, we examined their role in chemosignals-evoked neuronal activity, as well as in VNO-mediated innate behaviors. Using these multidisciplinary approaches, we demonstrate that the NPDs in rodents are not solely evolutionary remnant anatomical structures, but rather a key element in the biomechanical structure that allows constant pumping of chemosignals into the mammalian VNO and enables chemosignaling detection.

## Materials and methods

### Animals

Mature, sexually naïve, male C57BL/6 mice (Harlan Laboratories, Israel) were used in this study. Mice were maintained on a reverse 12/12 hours light/dark cycle, with food and water *ad libitum*. All experimental procedures were approved by the Institutional Animal Care and Use Committee of the Weizmann Institute of Science.

### Micro-CT

Mice were sacrificed and the oral openings of their NPDs were filled with radio opaque light curing hybrid composite with flowable viscosity (FLOWline, Heraeus Kulzer, Inc, IN, USA), in order to allow clear vision of the location and structure of the ducts in the micro-CT scan. The upper jaw of the mice was then removed and placed overnight in 4% paraformaldehyde fixative solution (PFA). Following fixation, samples were stained for 48 hours in Lugol solution (10g KI and 5g I_2_ in 100ml water) diluted 1:4 in DDW to generate an isotonic medium which minimizes the shrinkage of the soft tissue (Degenhardt et al 2010, Kelly 1961). Samples were then immobilized and sealed in a cylindrical holder made of polycarbonate. In order to avoid excessive tissue drying during the measurement, a small piece of wet cloth was placed at the bottom of the holder. This ensured water vapor saturated atmosphere around the sample for the whole duration of the experiment (about 30 hours). The holder was then firmly inserted into the sample support of the micro-CT instrument (Micro XCT-400, Xradia Ltd, California, USA). For the micro-CT scan, we set the X-ray source at 40KV and 200µA and took projection images with an objective having a nominal magnification of 0.5x. The scan included 6,000 such images taken with five seconds exposure time. No source filter was used. After volume reconstruction (done by the XRadia software which uses the Feldkamp algorithm for filtered back projection), we obtained final 3D images with 10µm resolution. Further image analysis was performed using Avizo software package (VSG Ltd, Bordeaux, France).

### Nasopalatine ducts blocking

Mice were randomly divided into experiment group (*blocked*) and sham operated group (*sham*). Animals were deeply anesthetized using Ketamine (100mg/kg) /Xylazine (23mg/kg), placed on their back, and their lower jaw was gently opened. A standard surgical cautery system (Gemini cautery kit, SouthPointe Surgical Supply Inc, Florida, USA) was used to block the oral entrance to the NPDs in the *blocked* group. Specifically, the heated tip of the cautery forceps (0.4mm in diameter) was placed at the entrance of each duct in the upper palate of the mouse, and cauterization was applied until adhesion of the tissue was visually observed (~500msec). In the *sham* group, the cautery forceps were placed on the upper palate just below the entrance to the ducts, and cauterization was applied as in the *blocked* group. Animals were monitored daily following the procedure and allowed 2-3 weeks to recover before the onset of experiments.

#### Confirmation of NPDs obstruction

For visual confirmation, Animals were anesthetized as described above. The NPDs openings where carefully examined under a binocular microscope (Nikon SMZ 745T) and photographed. For histological conformation, mice were sacrificed at the end of the experiment and their upper jaw was removed and placed in 4% PFA for a period of 7-days. Following fixation, the tissue was placed in 10% EDTA solution in room temperature for 10 days to allow decalcification (solution was changed every 3 days). The tissue was then washed in distilled water for two hours and in 50% ethanol for 30 minutes, before being embedded in paraffin. Coronal sections (7µm) of the complete palate and nasal cavity of each mouse were serially cut and mounted onto glass slides. The slices were stained using standard Hematoxylin-Eosin protocol (Burck 1973), and were examined to confirm closure of the ducts. Mice with two or more consecutive slices where the entrance to the ducts was not fully blocked were excluded from the analysis.

### Florescence dye assay

A 10µM rhodamine B solution (Sigma Laboratories) was freshly made at the beginning of each experiment week, and kept in 4°C, in dark condition, for a maximum of 7 days. Experimental mice from both groups (*blocked, sham*) were gently held in place and a total of 3µl of dye-mixture solution was gradually applied to their left nostril while allowing the mice to freely sniff the solution. Additional control group (*blank*) was comprised of *sham* mice that did not receive any stimulus, and used to quantify baseline autoflorescence levels in untreated VNO. Immediately after the dye-stimulus mixture was delivered, mice were euthanized, and their upper jaw was extracted and washed in 0.1M PBS solution. The upper palate was then removed, and the vomeronasal organ (VNO) was extracted bilaterally and washed with PBS. For measurement of florescence intensity, images of both side of each VNO were taken using a florescence stereomicroscope (Leica MZ FL III, Leica, Switzerland). Measurements of florescence were assessed using ImagePro Plus software (Media Cybernetics, Rockville, MD, USA). Mean optical density values were separately extracted for each side of each VNO, and then averaged to receive a single optical density value per mouse.

### Behavioral assays

#### Olfactory preference tests

Mice were individually housed for 1-2 weeks before initiation of behavioral trials. Prior to each experiment, pre-tests were conducted to exclude side preference in the testing apparatuses / home cage. At the beginning of each experiment day, animals were moved to the experiment room and allowed at least 1 hour to acclimate. For the odor preference assay, two applicators with cotton tips containing the different stimuli were attached to opposite walls of the home cage. On the first day of the experiment, mice were presented with one *“control stimulus”* (saline) and one *“social/neutral odor stimulus”* (200µl, male/female urine for social odor or banana/cinnamon for neutral odor); on the following day, mice were presented with one “*control stimulus”*, and the complementary *“social/neutral odor stimulus”*. Predator, vaginal secretion and saliva preference assays were conducted using a 3-chamber apparatus as previously described (Beny-Shefer et al 2017, Beny & Kimchi 2016, Karvat & Kimchi 2013, Zilkha et al 2017). Briefly, the apparatus is comprised of a polycarbonate box (70×24×29 cm) with partitions dividing the box into three chambers: a center chamber (15×24×29 cm) and two main chambers (25×24×29 cm). The partitions have retractable doorways (6.5×6.5 cm) allowing the animal to freely move between the chambers. Mice were allowed 10 minutes habituation to the setup, following which a social stimulus and a control stimulus were presented in the opposite main chambers. Mice were then allowed 10 minute to freely explore the apparatus. Olfactory investigation behavior was recorded using digital video cameras for later behavioral analysis. Social stimuli were as followed: for predator signals, soiled rat bedding was placed in a polycarbonate cup (5cm height X 7.5cm diameter). Saliva (100µl) and vaginal secretion (50µl) stimuli were presented on microscope slides attached to the chambers’ floor. The nature of the odor stimuli and the presentation sides were counter-balanced between mice. All tests were performed during the dark phase and under dim red light. Sniffing duration for each stimuli were analyzed using the Observer XT and Ethovision XT softwares (Noldus Information Technology, Wageningen, Netherlands Noldus). Mice with total sniffing time of less than 5% of overall experiment duration were excluded from the analysis. Absolute sniffing durations of each stimulus were calculated per mouse by subtracting time spent sniffing the control stimulus (e.g: *female exploration = duration sniffing female urine (sec) – duration sniffing saline (sec)*).

For urine stimuli, fresh urine was collected from 8-10 adult male or female C57BL/6J mice. Stimuli were kept in −80°C until use. Urine stimuli were diluted 1:1 by volume with saline. For control stimulus, standard saline solution was used. For predator stimuli, soiled rat bedding was collected from an adult Wistar rats cage, while the same amount of clean bedding was used as control stimulus. Saliva stimuli were collected from 15 adult male and 15 adult female C57BL/6J mice. Mice were anesthetized using Ketamine (100mg/kg) /Xylazine (23mg/kg) and exacerbated saliva secretion was induced via *Polocarpine* injection (0.025%, 100 µl, i.p.). Female saliva was diluted 2:3 by volume with saline, and standard saline solution was used as control stimulus. Vaginal secretions were collected from 7 adult female mice as previously described (McLean et al 2012, Scott et al 2015), with DDW used as control stimulus. For general odors, commercial cinnamon and banana odorant were used.

### Resident intruder assay

Intruder mice were sexually naive C57BL/6J females and males. A day prior to the experiment, female intruders were exposed to soiled male bedding in order to induce an estrous state. Resident male mice were introduced to the intruder in their home cage, and allowed to freely interact for 15 minutes. Social interaction was observed and recorded using digital video cameras, and analyzed offline using the Observer software (Noldus). The following behavioral parameters were measured: olfactory investigation, sexual behavior, aggression and locomotion activity.

### Food finding assay

Twenty-four hours prior to the experiment, food was removed from the home cage of the experimental mice and replaced with a small amount of food reward (one pine nut, ~0.1gr), in order to avoid food neophobia during behavioral tests. Before each trial, a single pine nut was randomly placed at the bottom of a large clean cage (20×35×18cm) covered with 2cm of bedding. Mice were then individually placed in the cage for five minutes and latency to discover the buried food was measured and compared between groups (*blocked, sham*).

### cFos induction following urine exposure

Urine was collected from 8-10 adult female mice in all stages of the estrous cycle. For control stimulus, double distilled water (DDW) was used. Animals were divided into three experimental groups: (1) *sham+DDW*, sham operated mice that were presented with 200μl of DDW; (2) *sham+urine*, sham operated mice that were presented with 200μl of female urine; (3) *blocked+urine*, mice with surgically blocked NPDs that were presented with 200μl of female urine. The night prior to the experiment, mice were individually placed in an empty cage with clean bedding. On the day of the experiment, stimulus (either urine or DDW) was placed on a small, round (~5cm in diameter), transparent and open Petri dish and positioned in the middle of the experimental cage. Mice were allowed to freely explore the stimulus for 15 minutes.

### Immunocytochemistry

One hour after stimulus presentation, mice were euthanized and perfused with cold 0.1M PBS followed by 4% PFA, as previously described (Scott et al 2015). The upper jaws of the mice were removed and examined to verify complete blocking of the ducts, as described above (see, *Confirmation of NPDs obstruction*). Brains were removed and post-fixed in 4% PFA for 48 hours. Using a vibratome (Leica Microsystems Inc.), brains were sliced into 30μm free-floating coronal sections. Sections were washed three times in 0.1M PBS and incubated for 30 minutes in a PBS/50% Methanol/0.32% HCL/1% H_2_O_2_ solution. After repeated washing in 0.1M PBS, slices were blocked using PBS/20% normal horse serum/0.2% Triton X-100 solution (1 hour), and incubated over night at 4°C in rabbit anti-cFos primary antibody solution (SC-52, 1:1,500, Santa Cruz Bio/technology, Santa Cruz, CA, USA). The following day, sections were again washed with 0.1M PBS and incubated in biotinylated goat anti-rabbit secondary antibody solution (1:200; Vector Laboratories, Burlingame, CA, USA) for 1 hour. Sections were then processed in ABC reagent (Vector Laboratories) for 1.5 hours and stained with diamino benzidine (DAB, Sigma Laboratories). cFos expression was assessed on both hemispheres of 5 sections per anatomical area. All labeled cell nuclei within the borders of the neuroanatomical nucleus of interest were counted using the ImagePro Plus software. The absolute number of labeled cells was counted and divided by area size in each slice to receive density values of number of cells per mm^2^. Density values from all five slices were then averaged for every mouse. This was done separately for each of the following anatomical regions of interest: anterior medial amygdala (aMeA), posterior medial amygdala (pMeA), anterior piriform cortex (aPir) and posterior piriform cortex (pPir), all as indicated in the mouse brain atlas (Paxinos. & Franklin. 2003).

### Statistical Analysis

All statistical analyses, unless stated otherwise, were performed by one-way or two-way ANOVA, followed by Fisher LSD post hoc comparisons as follows: Olfactory preference test was analyzed using two-way ANOVA with experimental group (*sham*/*blocked*) and stimulus (*male*/*female*) as main factors. For the florescence dye assay, latency to find the burried food, and social interactions in the resident intruder assay, groups (blocked, sham) were compared using the Mann-Whitney U test. Group data in the cFos counts was analyzed using one-way ANOVA (sham+DDW, sham+urine, blocked+urine). All statistical analyses were performed using the SPSS software (SPSS Inc., Chicago, USA), and STATISTICA software (StatSoft Inc, Tulsa, Okla). Mice scoring ±2STD away from group’s average were excluded from the analysis. Results are presented as mean±SEM, and the appropriate significant results are reported in detail when p<0.05.

## Results

### Micro-architecture of the NPDs presents continuous route between the nasal and oral cavities via the VNO

To establish the role of the NPDs in mammalian chemosignaling, we first examined whether they remain an open passageway connecting the oral cavity with the murine VNO. We detected the outer location of the NPDs openings in the upper palate of a mouse, on the border between the soft and the hard palate (Figure 1A,B). Then, we utilized a high-resolution micro-CT scanning technique with custom-designed methodology in order to reconstruct the complete 3-D morphological architecture of the nasal cavity and the nasopalatine ducts of mice (Figure 1 and supplementary Movies S1-3). The scans show that much like in reptiles, the murine NPDs constitute a direct passageway connecting the nasal cavity and the vomeronasal organ. The images show that the NPDs open at the posterior end of the VNO, engulf its bone-capsule and continue to the nasal cavity, while creating a clear route between the nasal cavity, the VNO and the mouth (Figure 1 and supplementary Movies S1-3). Specifically, the scans revealed a net of micro-tunnels (~100µm in diameter) which start at the nostrils and then branch into two main routs: one leads to the rear end of the nasal cavity, while the other leads to the VNO (Figure 1E). The tunnels reaching the VNO engulf it from all sides and connect directly to the NPDs, thus creating a continuous nostrils-VNO-mouth track (Figure 1C,D). In addition, unlike previous reports of a single anterior opening of the vomeronasal organ, we demonstrate here that another posterior opening in the VNO capsule is clearly visible in our CT images (Figure 1E). This opening connects directly to the observed network of micro-tunnels leading to the NPDs.

**Figure 1.**
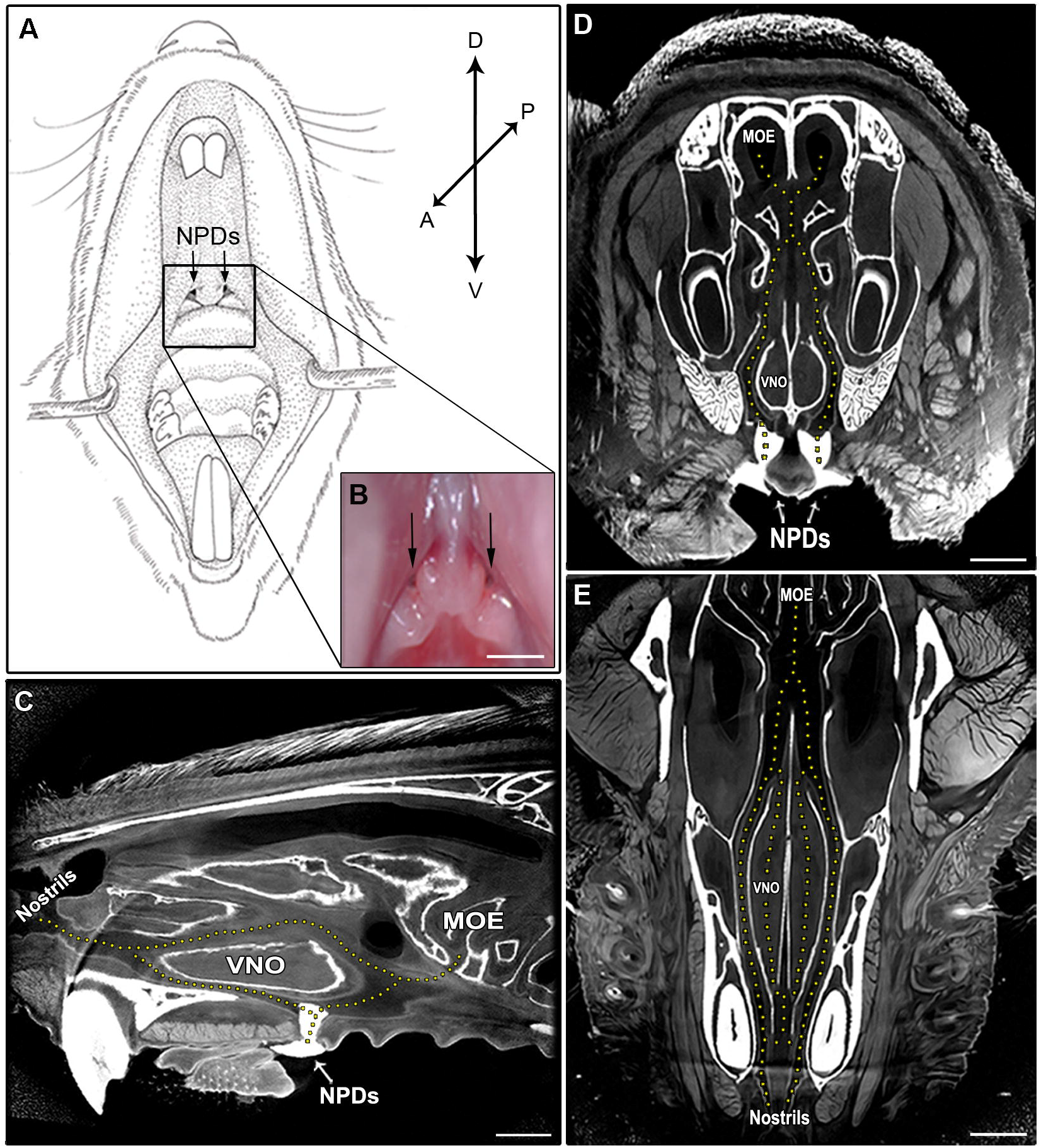
The mouse nasopalatine ducts facilitates substance flow through the VNO; (A) Schematic illustration of the oral cavity of an adult mouse. Location of the oral openings of the nasopalatine ducts is indicated by a black rectangle. (B) Image of the upper palate of a mouse. Arrows in the oral cavity indicate the two openings of the NPDs. (C,D,E) High-resolution micro computerized tomography (micro-CT) imaging of the mouse snout; (C) sagittal, (D) coronal and (E) transverse planes of a micro-CT scan (10µm resolution). Openings of the NPDs are filled with metallic high-contrast substance shown in white. The micro-CT scan revealed a complex network of pathways connecting the nasal cavity with the oral cavity and the VNO (indicated by dotted lines. see supplementary Movies S1-3). D-dorsal, V-ventral, A-anterior, P-posterior; VNO - vomeronasal organ; MOE - main olfactory epithelium; NPDs - nasopalatine ducts. Scale bar: 1mm.

### The NPDs are required for the pumping mechanism of the VNO

We then tested the hypothesis that the NPDs constitute a part of the active pumping mechanism that facilitates the flow of liquid-borne chemosignals to the receptor cell layer of the VNO (Eccles 1982, Meredith 1994). Considering the location of the ducts at the posterior part of the VNO, we speculated that they create a pathway for clearance of substance from the organ. To test this hypothesis we established a novel technique to obstruct the NPDs without damaging the VNO or the oral and nasal cavities. To do so, we used a standard cautery unit and applied it to the oral entrances of the NPDs found in the upper palate of adult male mice (Figure 1A,B), until adhesion of the local soft tissues was observed (*blocked* group) (Figure 2C,D). As control, we defined a *sham* group, where the cautery forceps were placed on the upper palate just below the openings, and cauterization was applied to the adjacent palate tissue as in the *blocked* group (Figure 2A,B).

**Figure 2.**
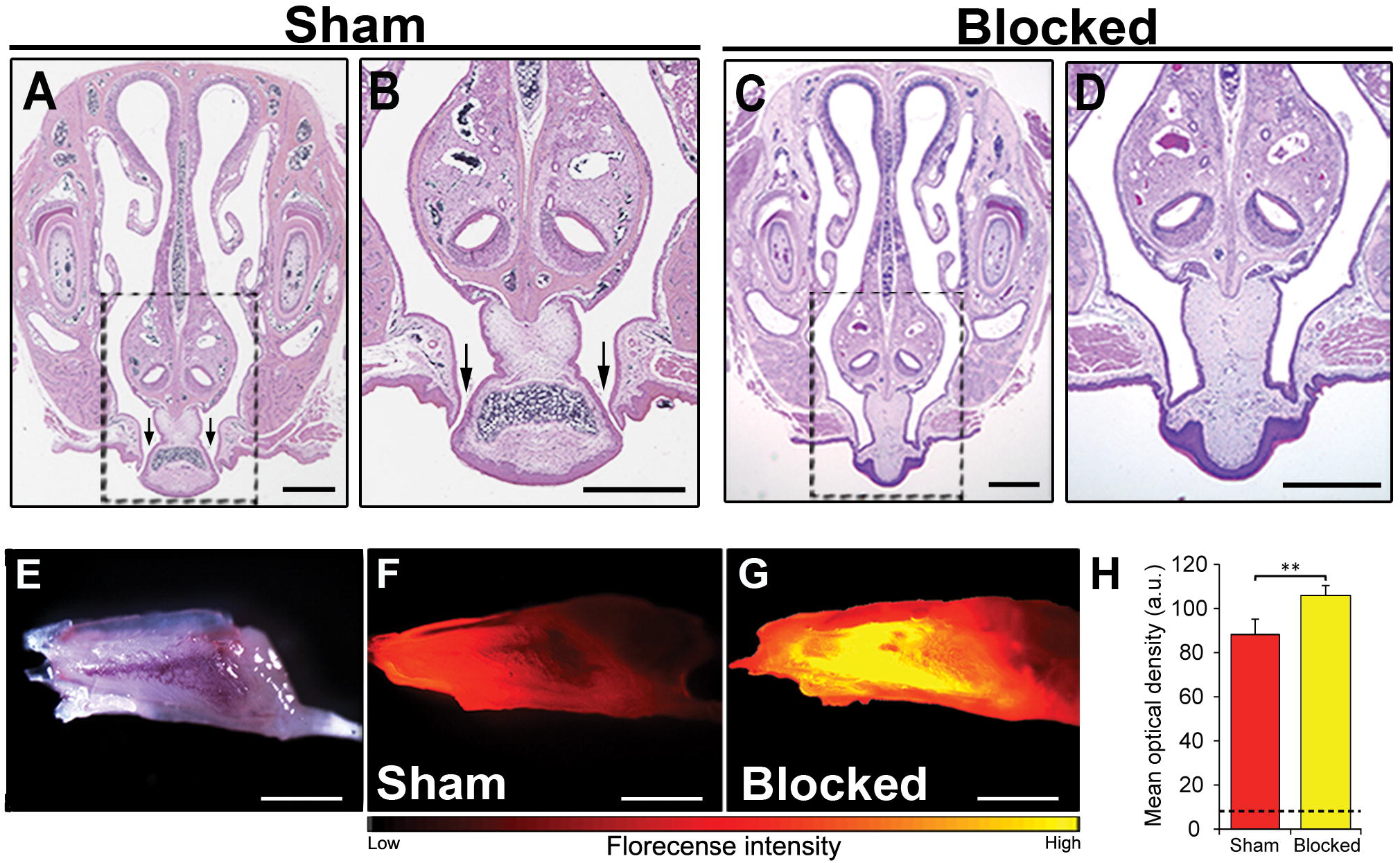
Obstruction of the NPDs leads to liquid accumulation in the VNO. (A,B,C,D) Coronal section through the snout and upper palate of a mouse with intact (A,B) or cauterized (C,D) NPDs, stained with standard HE staining. Rectangles in panels A,C are enlarged in panels B,D respectively. Black arrows indicate the oral openings of the NPDs. (E) Whole mount untreated VNO as seen under bright illumination. (F,G) Representative images of the VNO extracted from *sham* (F) and *blocked* (G) mice, following active nasal inhalation of rhodamine-tagged (red) liquid. Excess accumulating of liquid can be seen in the VNO of the *blocked* group. (H) Quantification of florescence signals in VNO following rhodamine treatment in *sham* (red) and *blocked* (yellow) groups. Dashed black line represent baseline mean optical density measured in control *sham* mice that did not receive any rhodamine (*blank*). Scale bar: 1mm. ** *p*<0.01. a.u. - arbitrary units.

To test whether such a procedure will indeed impair the VNO’s flow mechanism, we used a simple yet robust paradigm described in a study by Wisocky et al. (Wysocki et al 1980). We first allowed *blocked* and *sham* adult mice to sniff rhodamine-stained solution. We then measured the subsequent florescence levels in their VNO as an indication of the amount of substance to reach the lumen. In line with our hypothesis, the results show that all *blocked* mice presented an abnormal accumulation of dyed solution in their VNO (Figure 2E-H). Quantification of this signal revealed significantly higher levels of florescence in the VNO of *blocked* mice when compared to *sham* mice (n_sham_=7, n_blocked_=7, z=2.747, p=0.004; Figure 2H). This indicates that substances reaching the VNO are not properly cleared out in *blocked* mice, suggesting a malfunction in the pumping mechanism of the VNO.

### Obstructing the NPDs impairs chemosignaling-evoked neuronal activation

We next hypothesized that perturbation of the VNO’s pumping mechanism, via obstruction of the NPDs, can lead to deficits in VNO-mediated detection of chemosignals. Such impairment could be manifested in altered pheromone-evoked neuronal activity in brain regions involved in chemosignals processing. To directly test this notion, we first exposed male mice from both experiment groups (*blocked* and *sham*) to female urine (*blocked+urine* and *sham+urine*, respectively). An additional control group comprised of *sham* mice that were exposed only to distilled water (*sham+DDW*), as a measurement of cFos baseline activity. We measured neuronal activity levels in these groups by quantifying cFos immunoreactivity in the medial amygdala – a region known to be highly involved in the processing of chemosignals (Meredith & Westberry 2004, Petrulis 2013, Samuelsen & Meredith 2009a) and execution of social behaviors (Felix-Ortiz & Tye 2014, Shemesh et al 2016). We found that mice with *blocked* NPDs present significantly decreased neuronal activity levels in the anterior and posterior MeA when compared to *sham* mice, following active investigation of female urine. Importantly, this reduction was not observed in the piriform cortex, which was used as a control region. (aMeA, n=23, F_(2,20)_=30.323, p<0.001; post-hoc p<0.01 for *sham+urine* vs. *blocked+urine*; pMeA, n=24, F_(2,21)_=11.652, p<0.001; post-hoc p<0.05 for *sham+urine* vs. *blocked+urine*; aPir, n=25, F_(2,22)_=9.724, p<0.001; p=0.9 for *sham+urine* vs. *blocked+urine*; pPir, F_(2,22)_=11.9942, p<0.001; p=0.88 for *sham+urine* vs. *blocked+urine*; Figure 3).

**Figure 3.**
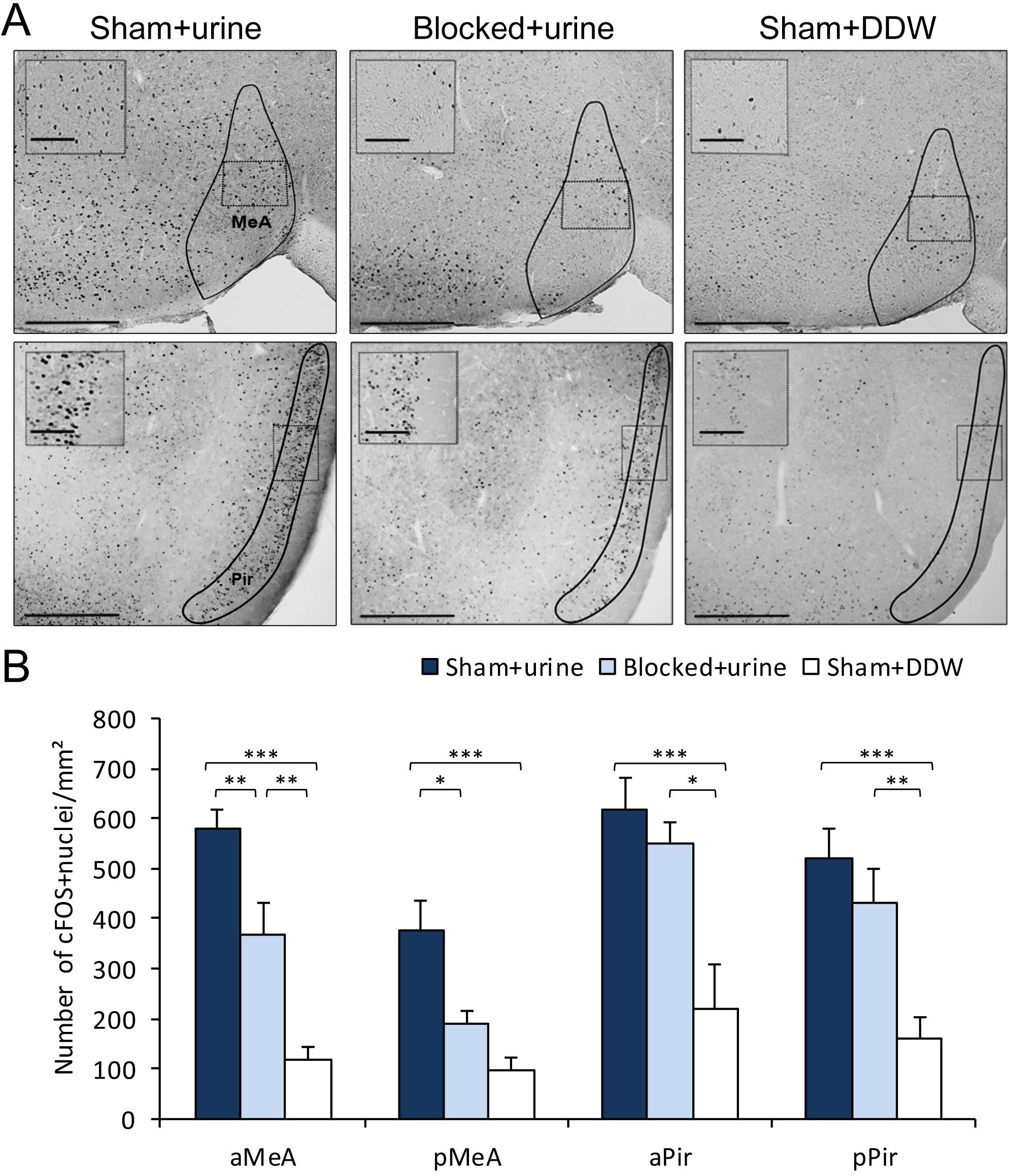
Blocking the NPDs results in significantly decreased neuronal activity in the vomeronasal system; (A) Representative images of cFos staining in coronal section of *sham+urine* (left panels), *blocked+urine* (middle panels) and *sham+DDW* (right panels) mice. Anatomical areas of interest are outlined in black. Insets depict areas outlined by dotted rectangles. (B) Quantification of cFos reactivity in secondary chemosignal processing brain regions. *Blocked* male mice exposed to female urine presented decreased neuronal activity when compared to *sham* mice. aMeA - anterior medial amygdala; pMeA – posterior medial amygdala; aPir - anterior piriform cortex; pPir - posterior piriform cortex. Scale bar: 500µm, inset: 100µm. * *p*<0.05, ** *p*<0.01, \*\*\**p*<0.001.

### NPDs are crucial for VNO-mediated social behaviors in male mice

Direct impairments in VNO function were repeatedly shown to induce alterations in chemosignaling-dependent innate behaviors such as social interactions and sexual preference (Ben-Shaul et al 2010, Chamero et al 2007, Dulac & Torello 2003, Kimchi et al 2007, Leypold et al 2002). To assess whether obstruction of the NPDs is in itself sufficient to induce similar deficits, we conducted a battery of well-established behavioral assays designed to examine chemosignaling-related behaviors. We first examined the effects of blocking the NPDs on the innate preference of male mice for female chemosignals (Bean et al 1986, Beny-Shefer et al 2017, Beny & Kimchi 2016). We exposed both *blocked* and *sham* male mice to either saline, female urine or male urine stimuli, which were presented on opposite sides of their home cage (Figures S3). We then tested the preference of each mouse by quantifying the duration it spent sniffing each stimulus. The results reveal that *sham* mice presented robust preference for female urine over male urine, while *blocked* mice exhibited no such preference (n_blocked_=7, n_sham_=9. F_stimulus(1,14)_=12.251; p=0.003; *sham* group: p<0.01 for sniffing female urine vs. sniffing male urine; *blocked* group: p=0.23 for the same comparison, figure 4A).

**Figure 4.**
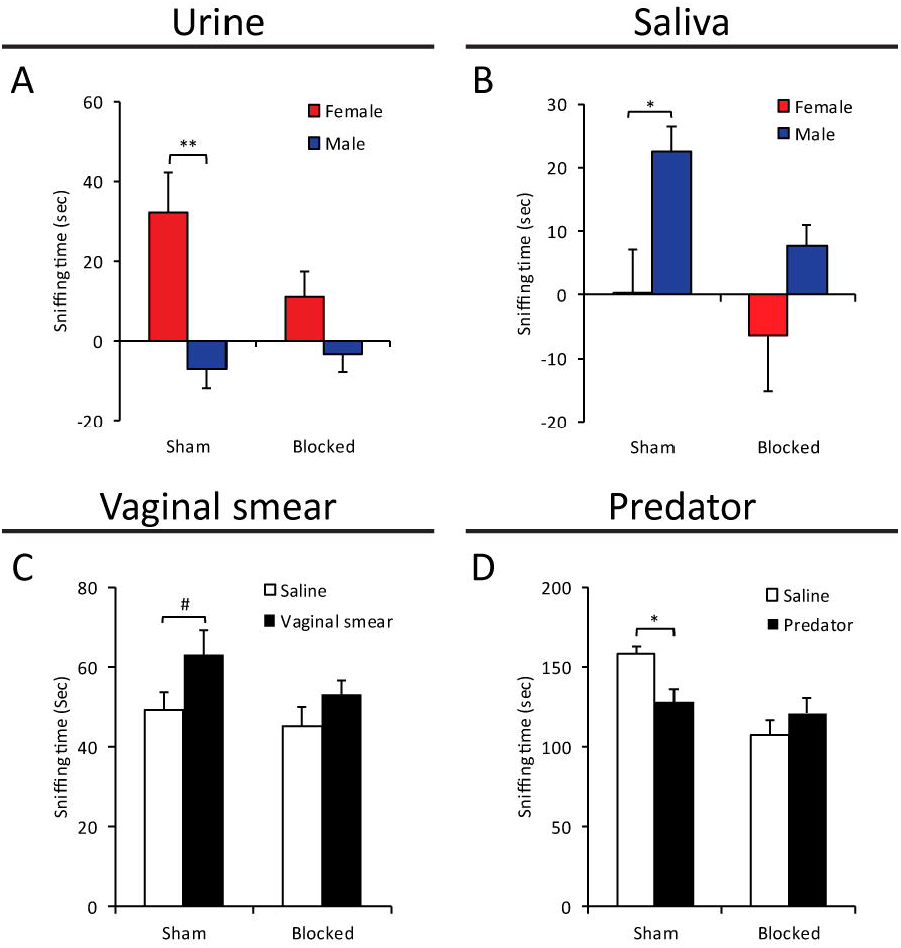
NPDs obstruction induces alterations in VNO-mediated responses. Mean duration of olfactory investigation in *sham* and *blocked* male mice presented with: (A) male/female conspecific urine, (B) male/female saliva, (C) female vaginal fluids, (D) predator bedding. Blocked mice presented significantly decreased preference for exploration of various chemosignals. #*p*=0.08, \**p*<0.05, \*\**p*<0.01, \*\*\**p*<0.001.

As chemosignals are found not only in urine, but also in other body secretions such as saliva (Gröschl 2009, Loebel et al 2000) and vaginal secretions (Bell et al 2013), we further tested for alteration in evoked behavioral responses to these stimuli in control and NPDs manipulated mice. We found that while *sham* mice preferred to explore saliva extracted from males over that female saliva, *blocked* mice exhibit no sex-specific preference (n_blocked_=12, n_sham_=9. F_stimulus(1,19)_=8.213; p<0.01; *sham* group: p<0.05 for *exploring male saliva vs. exploring female saliva*; *blocked* group: p=0.11 for the same comparison, Figure 4B). A similar effect was observed for exploration of vaginal secretion as *sham* mice showed a preference trend for exploring social chemosignal over a neutral stimulus (i.e. saline), while blocked mice did not show any clear preference. (n_blocked_=12, n_sham_=10. F_stimulus(1,20)_=4.652; p<0.05; *sham* group: p<0.08 for exploring vaginal secretions vs. saline; *blocked* group: p=0.253 for the same comparison; Figure 4C)

The VNO detects not only conspecific cues, but also danger interspecific signals such as molecules emitted from predators (known as karimones) (Papes et al 2010). We found that while control mice tend to avoid predator signals (rat-soiled bedding, in comparison to clean bedding), *blocked* mice did not behaviorally distinguish between the two stimuli, thus lacking the innate chemosignals-mediated predator avoidance response (n_blocked_=9, n_sham_=8. For exploration duration: F_stimulus X group(1,15)_=5.303; p=0.036; *sham* group: p<0.05 for sniffing predator bedding vs. clean bedding; *blocked* group: p=0.317 for the same comparison; Figure 4D).

Finally, to examine the effect of NPDs obstruction on free social interactions, we introduced mice from the different groups to both male and female intruders, and tested the duration of subsequent social exploration, sexual and aggressive behavior. We found that NPDs-*blocked* mice spend significantly more time investigating a female intruder compared to *sham* mice. In addition, the NPDs-*blocked* mice exhibited significantly lower locomotion activity in the presence of a female intruder (i.e. cage exploration), compared to *sham* mice (n_sham_=8, n_blocked_=12, for exploration duration: z=2.469, p<0.05, Figure 5A; for locomotion activity duration: z=-3.41, p<0.01, Figure 5C). No differences were found in the duration of sexual behavior (n_sham_=8, n_blocked_=12, z=0.99, p=0.316; Figure 5B). When conducting the same experiment but with male intruders, we found no differences in any parameter of social behavior between the mice with obstructed NPDs mice and control littermates (Figure S1).

**Figure 5.**
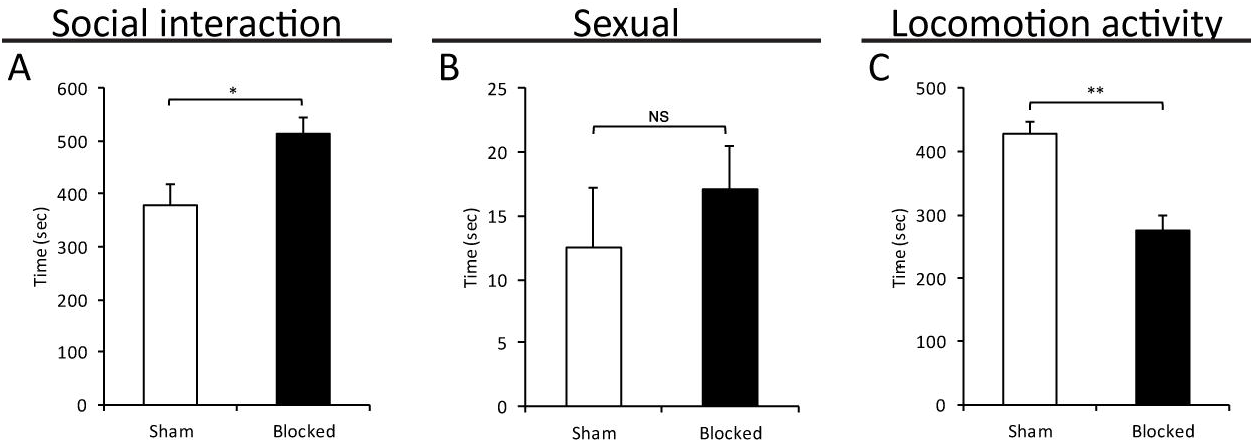
Obstruction of the NPDs induces specific impairments in VNO-mediated social behaviors. Significant alterations in social investigation (A) and in locomotion activity (C) in the presence of an intruder female, without detectable impairments in sexual behavior (B), in male mice with obstructed NPDs (*blocked*) vs. controls (*sham*). NS=not significant, \**p*<0.05, \*\**p*<0.01.

To control for the specificity of the observed behavioral responses to VNO-mediated behaviors, we conducted an additional set of behavioral assays designed to test for possible alterations in main-olfactory related behaviors (Fleming et al 2018, Vinograd et al 2017, Wilson et al 2006). First, we tested for changes in olfactory preference to non-social odors (cinnamon / banana), and found no differences in preference for these odors between the control and experiment groups (n_blocked_=9, n_sham_=9. For duration: F_treatment(1,16)_=0.014; p=0.99; banana: p=0.8 for *sham* vs. *blocked*; cinnamon: p=0.81 for the same comparison; Figure S2A,B). Next, we conducted a buried food-finding assay (Le Pichon et al 2009), where mice are placed in a cage with a hidden pine-nut that offers only olfactory cues for its location. No differences were found between groups in their latency to retrieve the concealed nut (n_sham_=14, n_blocked_=9, z=0.535, p=0.59. Figure S2C). Taken together, these results indicate no effect of obstructing the NPDs on main olfactory odor sensing.

## Discussion

The nasopalatine ducts are widely accepted as an integral part of the chemosignaling system in reptiles, creating the main route for pheromone transfer to the VNO. In mammals, and specifically in rodents, chemosignals are reported to reach the VNO through the nose by active sniffing (Shimp et al 2003, Wohrmann-Repenning 1980, Wohrmann-Repenning 1993). The question arises as to which role, if any, do the nasopalatine ducts play in the murine chemosignaling system. Our findings describe, for the first time, a functional role for the NPDs in facilitating substance flow to the mouse VNO. Based on high-resolution CT scans we report that the NPDs create a continuous route between the nostrils, the VNO and the mouth. Using *in-vivo* florescence tracing we demonstrated abnormal accumulation of fluids in the VNO following obstruction of the ducts, indicating that the mammalian NPDs constitute a route for substance clearing from the VNO.

Obstruction of the NPDs alone, with no perturbations of the VNO or the nasal pathway, resulted in prominent deficits in chemosignaling-evoked behavioral phenotypes. Specifically, NPDs-obstructed male mice lacked the innate preference towards female pheromones, a deficit that was observed in mice with surgically or genetically ablated VNO (Bean et al 1986, Kimchi et al 2007, Martinez-Garcia et al 2009, Nyby et al 1985, Pankevich et al 2004, Stowers et al 2002, Stowers & Logan 2010). Mice with ablated VNO also present an abnormally increased olfactory investigation behavior, a phenotype we have demonstrated as well in NPD obstructed mice. Moreover, NPDs-obstructed mice present impairments in sexual discrimination tasks (Bímová et al 2009, Byatt & Nyby 1986, Gröschl 2009), and showed no predator avoidance behavior (Blanchard et al 1998), indicating deficits in pheromone processing. However, NPDs-obstructed mice did not show any deficits in sexual behavior towards females, or in aggressive behavior towards males, unlike genetically or surgically ablated VNO mice (Dulac & Kimchi 2007, Leypold et al 2002, Stowers et al 2002).

The medial amygdala plays a crucial role in processing of social and predator signals detected by the VNO (Beny & Kimchi 2014, Bergan et al 2014, Chen & Hong 2018, Mohrhardt et al 2018). Here, we showed a significant reduction of neuronal activity (measured by cFos immunoreactivity) in the medial amygdala following NPDs obstruction, consistent with previous studies showing reductions in cFos expression following surgical removal of the VNO (Kondo et al 2003, Samuelsen & Meredith 2009b), further indicating that NPDs obstruction significantly impairs the function of the VNO.

It is well established that chemosignaling molecules enter the VNO lumen via an active pumping mechanism (Eccles 1982, Meredith 1994, Wysocki et al 1980). This repetitive pumping action requires both the active insertion and the active expulsion of chemosignals to/from the VNO. Considering this notion and our current results, we suggest that the murine NPDs are an essential component in this pump as they facilitate substance outflow from the VNO. This evacuation of fluids is crucial in order to enable the entrance of additional chemosignals into the VNO and the repetitive motion of the pump. Blocking the NPDs severely obstructs this continuous flow, leading the mechanism to impairments. Such obstruction will lead to deficits in pheromone processing and associated behavioral responses, as demonstrated by our behavioral and neuronal analysis. These results resemble some of the impairments observed in VNO ablated mice, although NPDs obstruction does not completely abolish VNO functioning, as some behavioral deficits found in VNO ablated mice were not found in our study. It should be noted, however, that while our findings indicate that the NPDs are a route for substance expulsion from the VNO, we cannot directly rule out that they might also constitute an inward route to the VNO for chemosignals collected through the oral cavity as is the case of reptiles. Nevertheless, our findings indicate a conserved functional role of the NPDs in murine chemosignaling, similar to reptiles.

## Acknowledgments

We thank C. Raanan and L. Friedman for their help in obtaining CT images, and the Kimchi lab members for their helpful comments on the manuscript.

---

**Supplementary Figure S1. Obstruction of the NPDs did not affect VNO-mediated social behaviors toward males.** Mean duration of (A) social investigation (B) aggression and (C) locomotion of male mice with obstructed NPDs (*blocked*) presented with an intruder male did not differ from control (*sham*) mice.

**Supplementary Figure S2. Male mice with NPDs obstruction present intact MOE-mediated olfactory investigation.** Preference for: (A) banana odor and (B) cinnamon odor. (C) Duration to find buried food in a buried food finding assay. No differences were found between *blocked* and *sham* groups.

**Supplementary Figure S3.** Experimental set-up of the urine olfactory preference assay. Two applicators containing the different stimuli (circled in red) were attached to opposite walls of the mouse home cage. Mice were allowed to freely investigate the two cotton tips for five minutes in each trial.

**Supplementary Movie 1.** Micro-CT 3D-reconstruction of the mouse snout: Coronal plane (from anterior to posterior).

**Supplementary Movie 2.** Micro-CT 3D-reconstruction of the mouse snout: Transverse plane (from ventral to dorsal).

**Supplementary Movie 3.** Micro-CT 3D-reconstruction of the mouse snout: Sagittal plane (from medial to lateral).

## References

Baum MJ, Kelliher KR. 2009. Complementary roles of the main and accessory olfactory systems in mammalian mate recognition. Annu Rev Physiol 71: 141–60

Bean NJ, Nyby J, Kerchner M, Dahinden Z. 1986. Hormonal regulation of chemosignal-stimulated precopulatory behaviors in male housemice (Mus musculus). Hormones and Behavior 20: 390–404

Bear Daniel M, Lassance J-M, Hoekstra Hopi E, Datta Sandeep R. 2016. The Evolving Neural and Genetic Architecture of Vertebrate Olfaction. Current Biology 26: R1039–R49

Bell MR, De Lorme KC, Figueira RJ, Kashy DA, Sisk CL. 2013. Adolescent gain in positive valence of a socially relevant stimulus: engagement of the mesocorticolimbic reward circuitry. European Journal of Neuroscience 37: 457–68

Ben-Shaul Y, Katz LC, Mooney R, Dulac C. 2010. In vivo vomeronasal stimulation reveals sensory encoding of conspecific and allospecific cues by the mouse accessory olfactory bulb. Proc Natl Acad Sci U S A 107: 5172–7

Beny-Shefer Y, Zilkha N, Lavi-Avnon Y, Bezalel N, Rogachev I, et al. 2017. Nucleus Accumbens Dopamine Signaling Regulates Sexual Preference for Females in Male Mice. Cell Reports 21: 3079–88

Beny Y, Kimchi T. 2014. Innate and learned aspects of pheromone-mediated social behaviours. Animal Behaviour 97: 301–11

Beny Y, Kimchi T. 2016. Conditioned odor aversion induces social anxiety towards females in wild-type and TrpC2 knockout male mice. Genes, Brain and Behavior 15: 722–32

Bergan JF, Ben-Shaul Y, Dulac C. 2014. Sex-specific processing of social cues in the medial amygdala. Elife 3: e02743

Bielsky IF, Young LJ. 2004. Oxytocin, vasopressin, and social recognition in mammals. Peptides 25: 1565–74

Bímová B, Albrecht T, Macholán M, Piálek J. 2009. Signalling components of the house mouse mate recognition system. Behavioural Processes 80: 20–27

Blanchard RJ, Hebert MA, Ferrari P, Palanza P, Figueira R, et al. 1998. Defensive behaviors in wild and laboratory (Swiss) mice: the mouse defense test battery. Physiology & Behavior 65: 201–09

Brennan PA, Zufall F. 2006. Pheromonal communication in vertebrates. Nature 444: 308–15

Burck HC. 1973. Histologische Technik. Georg Thieme Verlag Stuttgart: 110–11

Byatt S, Nyby J. 1986. Hormonal regulation of chemosignals of female mice that elicit ultrasonic vocalizations from males. Hormones and Behavior 20: 60–72

Chamero P, Marton TF, Logan DW, Flanagan K, Cruz JR, et al. 2007. Identification of protein pheromones that promote aggressive behaviour. Nature 450: 899–902

Chen P, Hong W. 2018. Neural Circuit Mechanisms of Social Behavior. Neuron 98: 16–30

Degenhardt K, Wright AC, Horng D, Padmanabhan A, Epstein JA. 2010. Rapid 3D phenotyping of cardiovascular development in mouse embryos by micro-CT with iodine staining. Circ Cardiovasc Imaging 3: 314–22

Dulac C, Kimchi T. 2007. Neural mechanisms underlying sex-specific behaviors in vertebrates. Current Opinion in Neurobiology 17: 675–83

Dulac C, Torello AT. 2003. Molecular detection of pheromone signals in mammals: from genes to behaviour. Nat Rev Neurosci 4: 551–62

Eccles R. 1982. Autonomic innervation of the vomeronasal organ of the cat. Physiol Behav 28: 1011–5

Felix-Ortiz AC, Tye KM. 2014. Amygdala inputs to the ventral hippocampus bidirectionally modulate social behavior. The Journal of Neuroscience 34: 586–95

Fleming G, Wright BA, Wilson DA. 2018. The Value of Homework: Exposure to Odors in the Home Cage Enhances Odor-Discrimination Learning in Mice. Chemical Senses 44: 135–43

Grinevich V, Stoop R. 2018. Interplay between Oxytocin and Sensory Systems in the Orchestration of Socio-Emotional Behaviors. Neuron 99: 887–904

Gröschl M. 2009. The physiological role of hormones in saliva. Bioessays 31: 843–52

Haga S, Hattori T, Sato T, Sato K, Matsuda S, et al. 2010. The male mouse pheromone ESP1 enhances female sexual receptive behaviour through a specific vomeronasal receptor. Nature 466: 118–22

Halpern M, Martinez-Marcos A. 2003. Structure and function of the vomeronasal system: an update. Progress in neurobiology 70: 245–318

Isogai Y, Si S, Pont-Lezica L, Tan T, Kapoor V, et al. 2011. Molecular organization of vomeronasal chemoreception. Nature 478: 241–5

Jacob S, Zelano B, Gungor A, Abbott D, Naclerio R, McClintock MK. 2000. Location and gross morphology of the nasopalatine duct in human adults. Arch Otolaryngol Head Neck Surg 126: 741–8

Karvat G, Kimchi T. 2013. Acetylcholine Elevation Relieves Cognitive Rigidity and Social Deficiency in a Mouse Model of Autism. Neuropsychopharmacology 39: 831

Kelly FC. 1961. Iodine in medicine and pharmacy since its discovery-1811-1961. Proc R Soc Med 54: 831–6

Keverne EB. 1999. The vomeronasal organ. Science 286: 716–20

Kimchi T, Xu J, Dulac C. 2007. A functional circuit underlying male sexual behaviour in the female mouse brain. Nature 448: 1009–14

Kondo Y, Sudo T, Tomihara K, Sakuma Y. 2003. Activation of accessory olfactory bulb neurons during copulatory behavior after deprivation of vomeronasal inputs in male rats. Brain Research 962: 232–6

Le Pichon CE, Valley MT, Polymenidou M, Chesler AT, Sagdullaev BT, et al. 2009. Olfactory behavior and physiology are disrupted in prion protein knockout mice. Nat Neurosci 12: 60–9

Leypold BG, Yu CR, Leinders-Zufall T, Kim MM, Zufall F, Axel R. 2002. Altered sexual and social behaviors in trp2 mutant mice. Proceedings of the National Academy of Sciences 99: 6376–81

Loebel D, Scaloni A, Paolini S, Fini C, Ferrara L, et al. 2000. Cloning, post-translational modifications, heterologous expression and ligand-binding of boar salivary lipocalin. Biochem. J 350: 369–79

Mackaysim A, Rose JD. 1986. Removal of the vomeronasal organ impairs lordosis in female hamsters - effect is reversed by Luteinizing-Hormone-Releasing hormone. Neuroendocrinology 42: 489–93

Marom K, Horesh N, Abu-Snieneh A, Dafni A, Paul R, et al. 2019. The Vomeronasal System Can Learn Novel Stimulus Response Pairings. Cell Reports 27: 676–84.e6

Martinez-Garcia F, Martinez-Ricos J, Agustin-Pavon C, Martinez-Hernandez J, Novejarque A, Lanuza E. 2009. Refining the dual olfactory hypothesis: pheromone reward and odour experience. Behavioural Brain Research 200: 277–86

McLean AC, Valenzuela N, Fai S, Bennett SA. 2012. Performing vaginal lavage, crystal violet staining, and vaginal cytological evaluation for mouse estrous cycle staging identification. J Vis Exp: e4389

Meredith M. 1991. Vomeronasal damage, not nasopalatine duct damage, produces mating-behavior deficits in male hamsters. Chem Senses 16: 155–67

Meredith M. 1994. Chronic recording of vomeronasal pump activation in awake behaving hamsters. Physiol Behav 56: 345–54

Meredith M, Westberry JM. 2004. Distinctive responses in the medial amygdala to same-species and different-species pheromones. Journal of Neuroscience 24: 5719–25

Mohrhardt J, Nagel M, Fleck D, Ben-Shaul Y, Spehr M. 2018. Signal Detection and Coding in the Accessory Olfactory System. Chemical Senses 43: 667–95

Nyby J, Kay E, Bean NJ, Dahinden Z, Kerchner M. 1985. Male-mouse (Mus-Musculus) attraction to airborne urinary odors of conspecifics and to food odors - effects of food-deprivation. Journal of Comparative Psychology 99: 479–90

Pankevich DE, Baum MJ, Cherry JA. 2004. Olfactory sex discrimination persists, whereas the preference for urinary odorants from estrous females disappears in male mice after vomeronasal organ removal. Journal of Neuroscience 24: 9451–7

Papes F, Logan DW, Stowers L. 2010. The vomeronasal organ mediates interspecies defensive behaviors through detection of protein pheromone homologs. Cell 141: 692–703

Paxinos. G, Franklin. KB. 2003. The mouse brain in stereotaxic coordinates Amsterdam: Elsevier Science and Technology Books.

Petrulis A. 2013. Chemosignals, hormones and mammalian reproduction. Hormones and behavior 63: 723–41

Samuelsen CL, Meredith M. 2009a. Categorization of biologically relevant chemical signals in the medial amygdala. Brain Research 1263: 33–42

Samuelsen CL, Meredith M. 2009b. The vomeronasal organ is required for the male mouse medial amygdala response to chemical-communication signals, as assessed by immediate early gene expression. Neuroscience 164: 1468–76

Scott N, Prigge M, Yizhar O, Kimchi T. 2015. A sexually dimorphic hypothalamic circuit controls maternal care and oxytocin secretion. Nature 525: 519–22

Shemesh Y, Forkosh O, Mahn M, Anpilov S, Sztainberg Y, et al. 2016. Ucn3 and CRF-R2 in the medial amygdala regulate complex social dynamics. Nature Neuroscience 19: 1489

Shimp KL, Bratnagar KP, Bonar CJ, Smith TD. 2003. Ontogeny of the nasopalatine duct in primates. Anat Rec Part A 274A: 862–69

Stahlbaum CC, Houpt KA. 1989. The role of the Flehmen response in the behavioral repertoire of the stallion. Physiol Behav 45: 1207–14

Stowers L, Holy TE, Meister M, Dulac C, Koentges G. 2002. Loss of sex discrimination and male-male aggression in mice deficient for TRP2. Science 295: 1493–500

Stowers L, Logan DW. 2010. Olfactory mechanisms of stereotyped behavior: on the scent of specialized circuits. Current Opinion in Neurobiology 20: 274–80

Vaccarezza OL, Sepich LN, Tramezzani JH. 1981. The vomeronasal organ of the rat. J Anat 132: 167–85

Vinograd A, Livneh Y, Mizrahi A. 2017. History-Dependent Odor Processing in the Mouse Olfactory Bulb. The Journal of Neuroscience 37: 12018–30

Wilson DA, Stevenson RJ, Stevenson RJ. 2006. Learning to smell: olfactory perception from neurobiology to behavior. JHU Press.

Wohrmann-Repenning A. 1980. The relationship between jacobsons organ and the oral cavity in a rodent. Zool Anz 204: 391–99

Wohrmann-Repenning A. 1993. The vomeronasal complex - a dual sensory system for olfaction and taste. Zool. Jb. Anat 123: 337–45

Wysocki CJ, Wellington JL, Beauchamp GK. 1980. Access of urinary nonvolatiles to the mammalian vomeronasal organ. Science 207: 781–83

Zilkha N, Kuperman Y, Kimchi T. 2017. High-fat diet exacerbates cognitive rigidity and social deficiency in the BTBR mouse model of autism. Neuroscience 345: 142–54

