## Supplementary figures and images for "The nasopalatine ducts of the mouse conserve a functional role in pheromone signaling"

### Supplemental figure 1

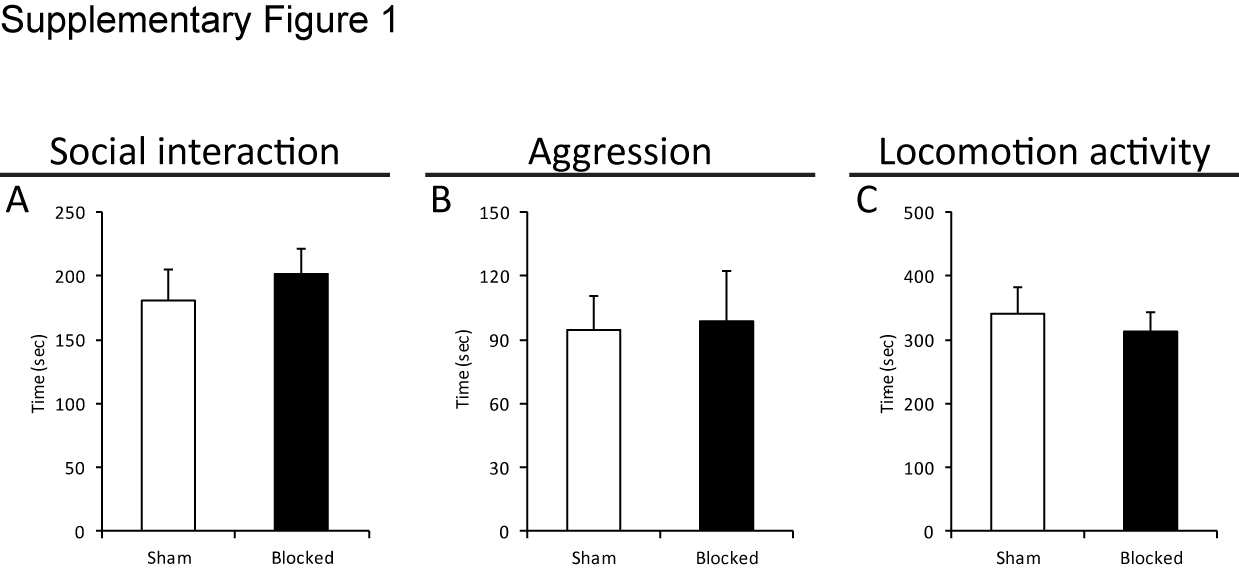

### Supplemental figure 2

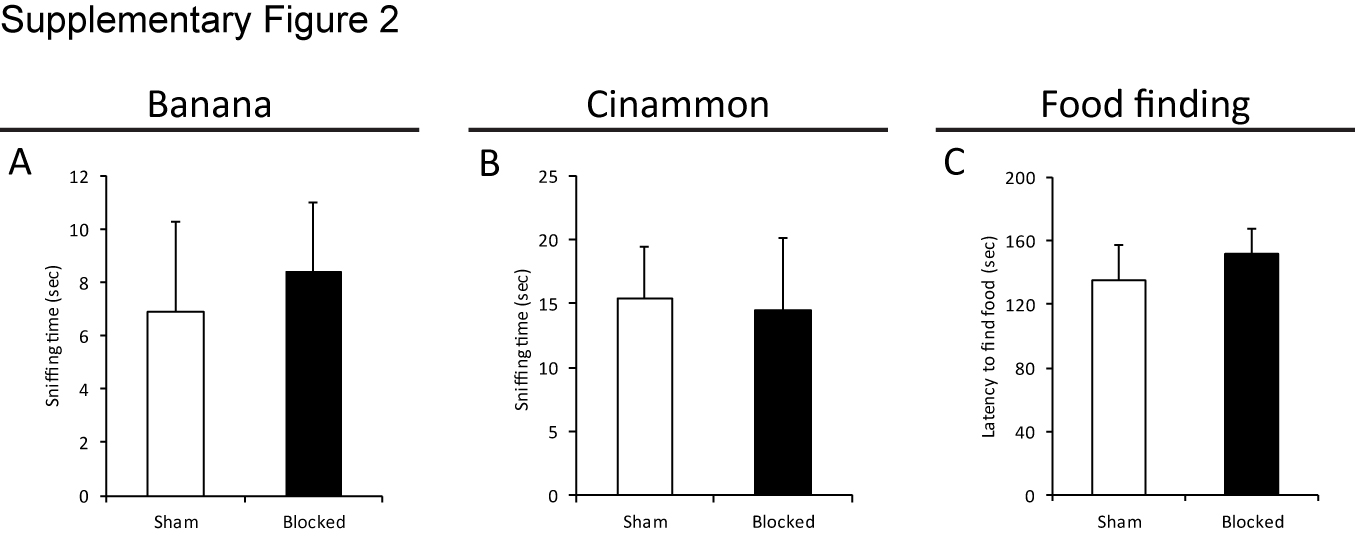

### Supplemental figure 3

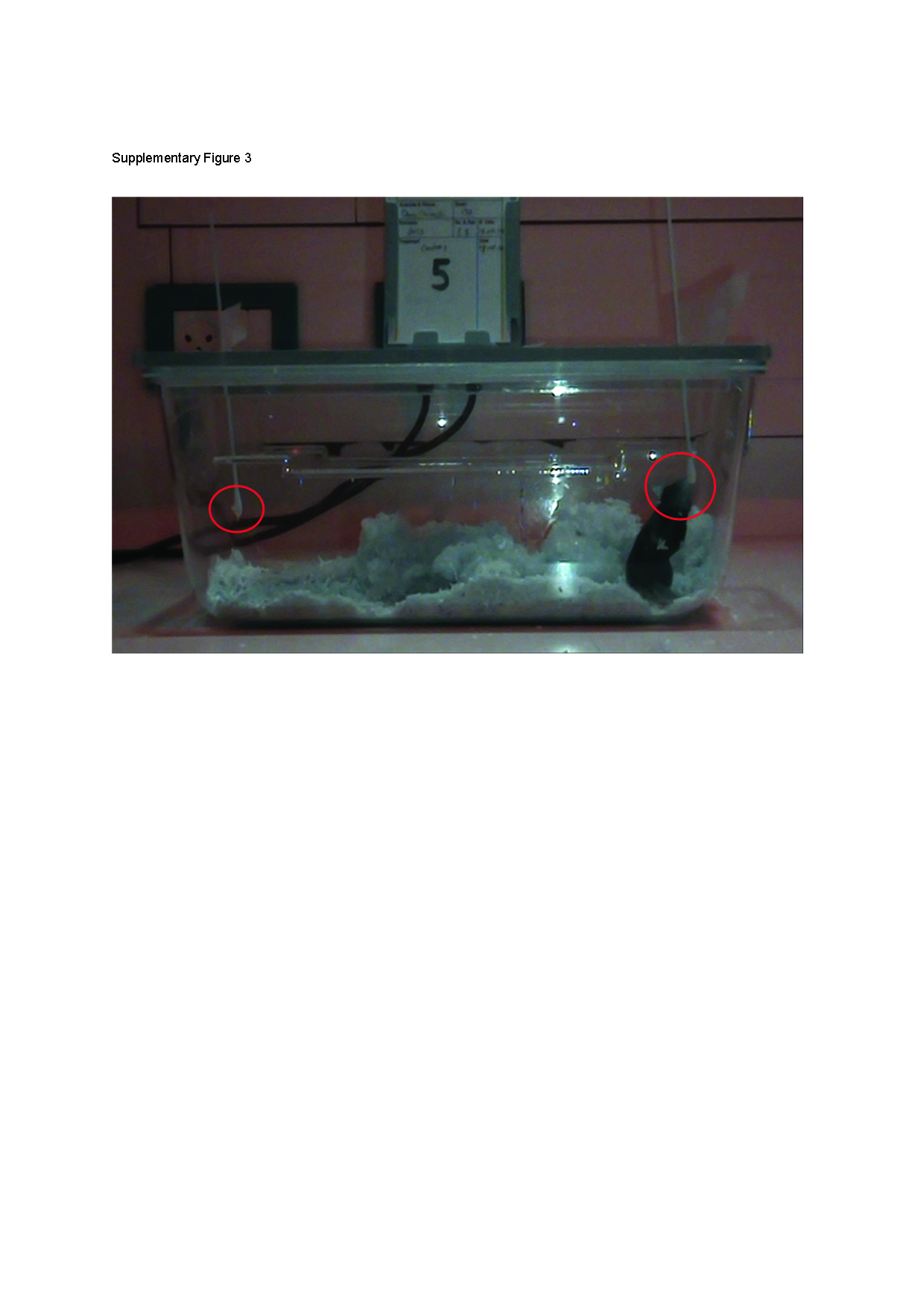
